# Strong purifying selection in haploid tissue-specific genes of Scots pine supports the masking theory

**DOI:** 10.1101/2023.04.11.536341

**Authors:** Sandra Cervantes, Robert Kesälahti, Timo A. Kumpula, Tiina M. Mattila, Heikki Helanterä, Tanja Pyhäjärvi

## Abstract

The masking theory states that genes expressed in haploid stage will be under more efficient selection. In contrast, selection will be less efficient in genes expressed in diploid stage, where the fitness effects of recessive deleterious or beneficial mutations can be hidden from selection in heterozygous form. This difference can influence several evolutionary processes such as maintenance of genetic variation, adaptation rate, and genetic load. Masking theory expectations have been confirmed in single-cell haploid and diploid organisms. However, in multicellular organisms, such as plants, the effects of haploid selection are not clear-cut. In plants, the great majority of studies indicating haploid selection have been carried out using male haploid tissues in angiosperms. Hence, evidence in these systems is confounded with the effects of sexual selection and intra-specific competition. Evidence from other plant groups is scarce and results show no support for the masking theory.

Here we have used a gymnosperm Scots pine megagametophyte, a maternally-derived seed haploid tissue, and four diploid tissues to test the strength of purifying selection on a set of genes with tissue-specific expression. By using targeted resequencing data of those genes, we obtained estimates of genetic diversity, the site frequency spectrum of 0-fold and 4-fold sites, and inferred the distribution of fitness effects (DFE) of new mutations in haploid and diploid tissue-specific genes. Our results show that purifying selection is stronger for tissue-specific genes expressed in the haploid megagametophyte tissue, and that this signal of strong selection is not an artifact driven by high expression levels

## Introduction

The masking theory predicts that the efficacy of selection is stronger in haploid genomes (Kondrashov and Crow 1991) in comparison to diploid, because the number of chromosomal copies in a genome directly affects the efficacy of selection. For genes expressed on diploid genomes, any level of dominance (*h*) other than 0.5 (additivity) implies that the fitness effect of one allele will be partially or totally masked by the other allele. Consequently, both deleterious and beneficial mutations can be less affected by selection and hide in a heterozygous state. In contrast, genes expressed in haploid genomes will be readily exposed to selection due to the lack of masking (Crow and Kimura 1965; Kondrashov and Crow 1991). This difference has important evolutionary consequences as selection acting differently in haploid and diploid genomes will affect the spread and fixation of new mutations and influence genetic load, genetic variation, and adaptation rate (Szövényi et al. 2013; Immler and Otto 2018).

Most of the empirical evidence supporting haploid selection has been obtained through yeast experimental evolution studies (Otto and Gerstein 2008; Gerstein et al. 2011), where haploid organisms adapt faster and have better fitness when exposed to environmental changes, e.g., nutrient limitation (Otto and Gerstein 2008 and references therein, Mable and Otto 1998). However, expectations of masking theory can be extended to multicellular organisms with alternation of generations between haploid and diploid phases, such as plants (Immler 2019; Beaudry et al. 2020). In these cases, selection in the haploid stage can help, for example, to reduce the burden of recessive deleterious mutations carried in the diploid stage, or it could lead to the evolution of heteromorphic life cycles such as ecological niche differentiation at haploid and diploid stages, which will maximize resource exploitation (Mable and Otto 1998).

Research on haploid selection in plants has been conducted with angiosperms, particularly in genera within the Brassicaceae family, e.g. Arabidopsis and Capsella (Arunkumar et al. 2013; Gossmann et al. 2014; Gutiérrez-Valencia et al. 2022), where the female gametophytic stage is very reduced and dissection of female structures can be technically difficult (Beaudry et al. 2020, but see Gossman et al. 2014 and Gutierrez-Valencia et al. 2021). Therefore, most of these studies have compared male gametophytes, typically pollen-expressed genes, to genes expressed in sporophytic tissues. Results of these studies have shown that selection on pollen (haploid) expressed genes is stronger relative to selection on sporophyte (diploid) expressed genes (Arunkumar et al. 2013; Gossmann et al. 2014). Studies in other angiosperm systems have shown similar results. In *Silene latifolia* and Rumex species, increased efficacy of purifying selection at the haploid stage allows the purging of deleterious alleles of Y-linked genes with expression at the male gametophytic stage, which in turn slows down the degeneration of their Y chromosome (Chibalina and Filatov 2011; Sandler et al. 2018).

However, it is argued that in angiosperms the increased selective pressure observed in haploid-stage expressed genes carries the confounding effects of pollen competition (Moore and Pannell 2011). The general assumption is that selection has a stronger effect on male haploid stages than on female stages, in part because in plants the pollen is released to an exterior heterogeneous environment, which implies exposure to varying degrees of environmental selective pressures. In comparison, plant female reproductive structures, gametophyte and eggs, remain sheltered inside the sporophyte (Immler and Otto 2018; Sandler et al. 2018). In addition, most of the female eggs will be fertilized, while only a small number of male gametes carried by pollen will be able to participate in the fertilization (Haldane 1924, Immler 2019). For example, Arunkumar et al. (2013) acknowledged that it is difficult to disentangle the signal of haploid selection from the signal of sexual selection in pollen of *Capsella grandiflora*.

Moreover, current knowledge on haploid selection mostly represents the evolutionary dynamic in angiosperms and does not represent the variety of length of the haploid stage in other plant groups such as mosses or gymnosperms. To our knowledge there is only one study in a non-angiosperm species (*Funaria hygrometrica*) where explicit testing of haploid selection has been approached (Szövényi et al. 2013). In *F. hygrometrica*, a moss with an extended haploid phase and short diploid stage, Szövényi et al. 2013 found that haploid expressed genes have higher sequence variation due to relaxed selection. This finding contradicts the expectations of the masking theory. Szövényi et al. 2013 argue that confounding effects of the evolutionary dynamics of gene expression per se could be the reason for inefficient purifying selection. Level and breadth of expression are both known determinants of protein evolutionary rate, and genes with expression on a higher number of tissues (broad breadth of expression) and genes with high level of expression are under tighter selective constraints (e.g., stronger purifying selection) (Duret and Mouchiroud 2000; Zhang and Yang 2015). Therefore, the low level and narrow breadth of expression of tissue-specific genes relax the selective pressure over them rendering haploid selection insufficient to purge putative deleterious variation. Similar results have been observed in sperm-expressed genes of *Capsella grandiflora* and *Arabidopsis thaliana*, where low level of expression has been suggested as the explanation for relaxed selection (Arunkumar et al. 2013; Gossmann et al. 2014).

Here we used *Pinus sylvestris*, a gymnosperm with high realized outcrossing level, large population size and low level of genetic structure, as a study model (Pyhäjärvi et al. 2020; Tyrmi et al. 2020; Hall et al. 2021). *P. sylvestris* has a haploid megagametophyte stage of approximately two years. The megagametophyte is functionally homologous to the endosperm in the seed of angiosperms. However, the origin of the megagametophytic tissue is completely different in comparison to angiosperms. This multicellular structure originates from the meiosis of the megaspore mother-cell, which develops in the megasporangia of the ovuliferous scales on the female strobilus. Hence, unlike the endosperm, the megagametophytic tissue does not undergo fertilization, and only represents the maternal genotype (Williams 2009). Beside this, the megagametophyte is a metabolically and transcriptionally active tissue (Vuosku et al. 2009; Cervantes et al. 2021) and does not carry the confounding effects of haploid male-tissue competition observed in angiosperms.

Here we have looked at the effect of purifying selection on the genetic diversity (π), the folded site frequency spectrum (fSFS), and the distribution of fitness effects (DFE) of genes expressed in haploid (megagametophyte) and diploid (embryo, vegetative bud, needle, and phloem) tissues of *P. sylvestris*. We chose to use tissue-specific genes as pleiotropic constrains arising from broad tissue expression could cause a confounding signal of purifying selection (Huber et al. 2017). Also, the use of tissue-specific genes allows a more reliable comparison across gene categories within species. Beside this, to avoid confounding effects due to demography, all samples come from a single population and our study is limited to within-species analyses. We expect that genes specific to the haploid stage in predominately diploid organisms, should display a similar response to selection as laid out by theoretical and empirical expectations of haploid selection in single-cell organisms. Hence, we expect to observe lower levels of genetic diversity and lower values of the four-fold to zero-fold pairwise nucleotide diversity ratio (π_0_/π_4_) for genes expressed in the haploid stage due to more effective purifying selection (Charlesworth et al. 1993; Charlesworth 1994).

Additionally, when efficient, purifying selection draws deleterious mutations to low frequencies. Thus, we also expect to observe a skew toward rare alleles in fSFS of haploid tissue-specific genes and a strong signal of purifying selection in haploid tissue-specific genes based on estimates of the DFE.

## Results

We base our study on genetic diversity and expression patterns of tissue-specifically expressed genes in four diploid and one haploid tissue. To assess the influence of purifying selection on the nucleotide diversity of the tissue-specifically expressed genes we genotyped 20 megagametophytes from unrelated trees of a single population using exome capture (Kesälahti R. unpublished data). We identify tissue-specific genes based on a previousl study (Cervantes et al. 2021). Analyses were done in six datasets, one for each tissue-specific set of genes, plus a dataset used as reference point which included all variable sites (hereafter refer as all-sites dataset).

### Genetic diversity level of genes with tissue-specific expression patterns

To identify the effects of purifying selection across genes with varying expression patterns, we first estimated nucleotide diversity at 0-fold and 4-fold sites (π_0-fold_, π_4-fold_) and their ratios per gene. Purifying selection reduces molecular genetic diversity due to background selection around the selected sites (Charlesworth et al. 1993; Charlesworth 1994) and reduces the π_0-fold_/π_4-fold_ ratio. In general, π and π_0_/π_4_ ratio values obtained in this study are within the range of genetic diversity known for *P. sylvestris* (Grivet et al. 2017; Pyhäjärvi et al. 2020). However, our results show no differences in the levels of neutral diversity (π_4-fold_) or ratios (Table 1, one-way ANOVA for five tissues, p=0.182) among tissues. Nevertheless, we did observe a tendency towards lower values of π_0-fold_/π_4-fold_ in needle and megagametophyte

**Table 1.** Summary of average pairwise nucleotide diversity per gene (π) for 0-fold and 4-fold sites and their ratios. All estimates were obtained from 1000 bootstrap of the mean. Estimates are provided for tissue-specifically expressed genes (tau score 0.8 to 1) across the five tissues and to all genes. SE = standard errors.

| | $\pi_{0\text{-fold}}$ (SE) | $\pi_{4\text{-fold}}$ (SE) | $\pi_{0\text{-fold}}/\pi_{4\text{-fold}}$ (SE) |
| --- | --- | --- | --- |
| Bud | 0.0009 (0.0001) | 0.0029 (0.0004) | 0.31 (0.05) |
| Embryo | 0.0014 (0.0002) | 0.0037 (0.0005) | 0.39 (0.07) |
| Megagametophyte | 0.0009 (0.0001) | 0.0032 (0.0004) | 0.29(0.06) |
| Needle | 0.0005 (6.85e-05) | 0.0021 (0.0003) | 0.25 (0.05) |
| Phloem | 0.0007 (0.0001) | 0.0025 (0.0004) | 0.31 (0.06) |
| All-genes | 0.0008 (4.65e-05) | 0.0028 (0.0002) | 0.30 (0.02) |

### Distribution of fitness effects of new mutations (DFE)

π estimates and their ratios are not optimal signals of purifying selection because they do not fully consider the effect of purifying selection on different classes of allele frequencies. There are, however, more sensitive methods to specifically estimate the strength of purifying selection based on site frequency spectra (SFS) that rely on the differences of observed and expected allele frequencies.

The DFE-alpha is a method for estimating the expected fitness effects of new mutations entering the population. It is based on the concept that the fitness effect of a mutation is one of the determinants of the frequency it will be found in the population (Eyre-Walker and Keightley 2007; Peter D Keightley and Eyre-Walker 2007; Keightley and Eyre-Walker 2010). Briefly, the DFE-alpha uses the observed amount of putative neutral diversity (4-fold) to estimate the population mutation rate and to account for the effects of demographic history. Using this information and the number of sites available for mutations to occur, the program can infer the amount of amino acid changing mutations that could have entered the population and compares it to the observed number of 0-fold sites. Considering all these components DFE-alpha allows the inference of the strength of purifying selection.

To estimate the DFE of different tissue-specifically expressed genes and all sites, we obtained fSFSs for 0-fold and 4-fold positions (Figure 1). A visual inspection of the fSFS shows that in the megagametophyte-specifically expressed genes, 4-fold sites had proportionately more singletons than 0-fold sites in contrast to all other site categories, and all other tissue-specific genes. Additionally, the difference between the proportion of 4-fold sites to 0-fold sites at the invariant category is larger in the megagametophyte compared to other tissue-specific expressed genes.

**Figure 1.**
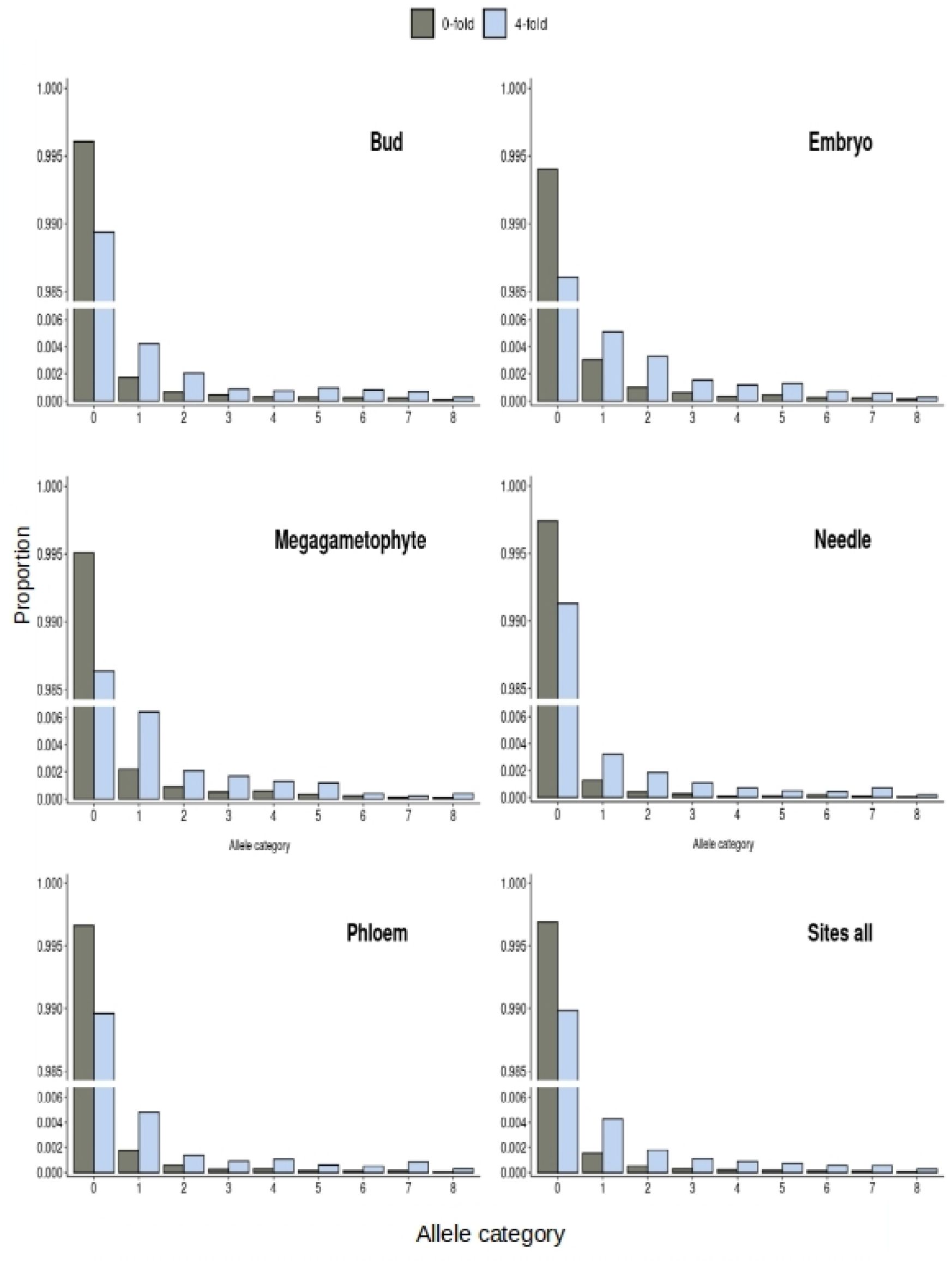
Folded site frequency spectrum for 0-fold and 4-fold sites at tissue-specific genes. The fSFS were obtained from a down-sampling of the observed data. The same data was also used as part of the input fSFSs to estimate the DFE (see Material and Methods section). y axis break was introduced using ggbreak (Xu et al. 2021).

To obtain the DFE of 0-fold sites of the five tissue-specific genes sets and the all sites, we used DFE-alpha 2.16 with a two-epoch model (Peter D Keightley and Eyre-Walker 2007). Further, all nucleotide diversity data came from a single population to avoid further confounding effects due to population structure. Based on visual inspection of the DFEs, tissue-specific genes had a distinct DFE from the all-sites data set which represents the genome-wide DFE (Figure 2). However, the tissue-specific genes did not have a consistent deviation from the genome-wide pattern. For example, the bud tissue-specific genes have a high proportion of mutations in the category N_e_s>100, which is an indicator of a stronger purifying selection, whereas phloem has fewer mutations in this category in comparison to the genome-wide average (Figure 2). Other differences among tissues, in comparison to all sites, are evident in all DFE classes as shown in Figure 2.

**Figure 2.**
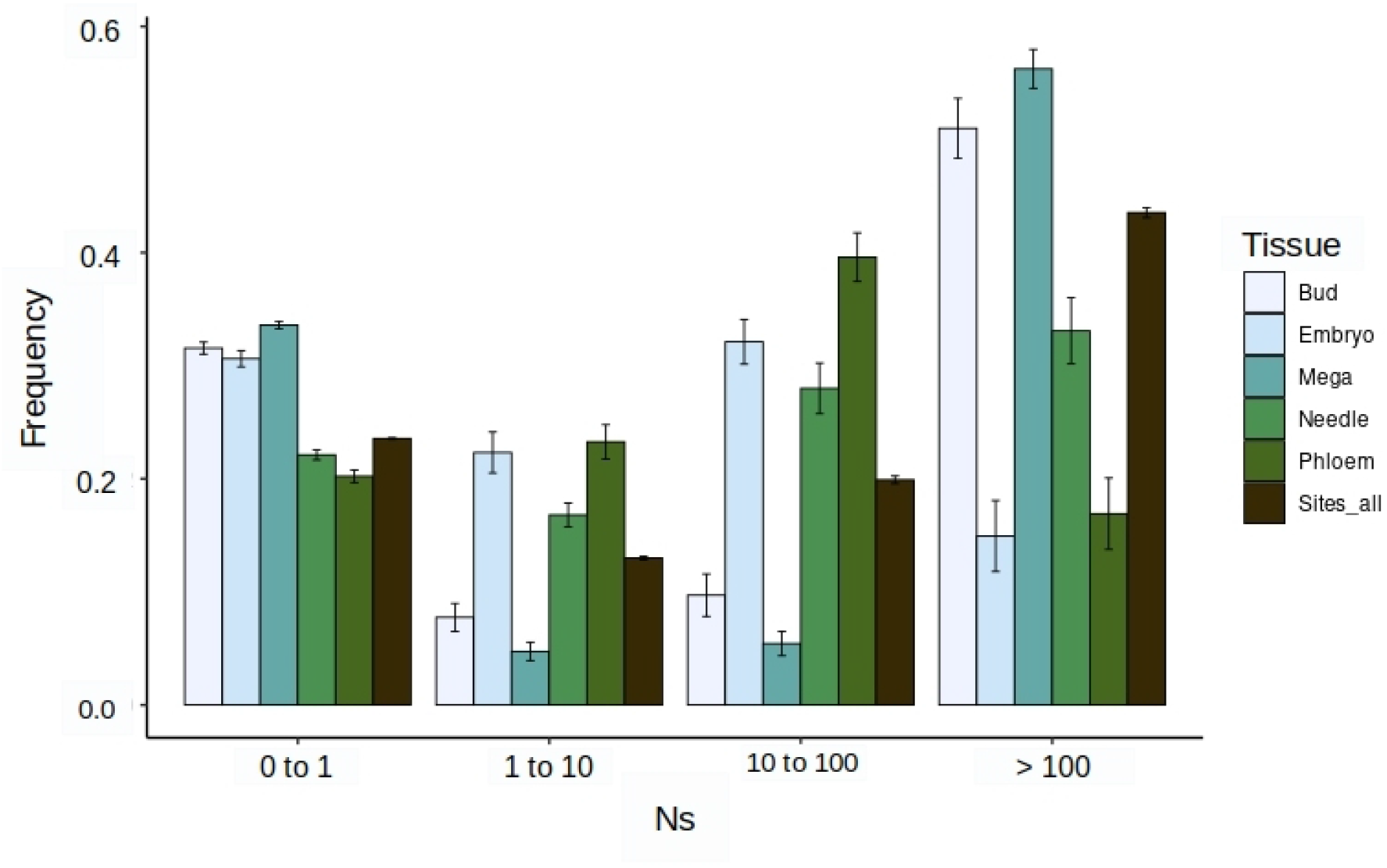
Distribution of fitness effects of tissue-specific genes (Tau score ≥ 0.8) for the five tissues and sites-all data sets. Inferences are based in 200 resamplings of the fSFS. Bars represent standard errors.

Interestingly, megagametophyte tissue-specific genes have the highest proportion of sites in the class N_e_s>100, indicating a large proportion of sites under strong purifying selection, but also display the higher proportion of neutral or nearly-neutral mutations (0<N_e_s<1 category). Overall, our results show that each tissue-specific set of genes has a distinct DFE, which is not surprising considering that different tissues are under different selective pressures according to their functionality, and in some cases depending on the developmental stage of the organ or tissue where they are expressed.

The DFE visualization is usually divided in four different classes according to scaled strength of selection (Figure 2). Classes are inferred from the β parameter (shape parameter of the gamma distribution) and the mean selective effect of a new mutation (Es) estimate (scale parameter of the gamma distribution). Low values of the β parameter (β → 0) are indicative of a highly leptokurtic gamma distribution (L-shaped gamma) with most of the mutations being either nearly neutral (low fitness effect) or strongly deleterious (high fitness effect) (Peter D. Keightley and Eyre-Walker 2007; Brevet and Lartillot 2021). To estimate how different the DFE of the megagametophyte tissue-specific genes is compared to the other tissues we inspected the distribution of the β parameter, instead of calculating confidence intervals over the different classes of strength of selection (Nes) as usually done. A distribution of the β parameter values obtained from 200 permuted DFE inferences is shown in Figure 3, where megagametophyte tissue-specific genes have the lowest values of the β parameter and their distribution only overlaps with that of the bud tissue-specific genes with the overlap accounting for approximately 7% of the values.

**Figure 3.**
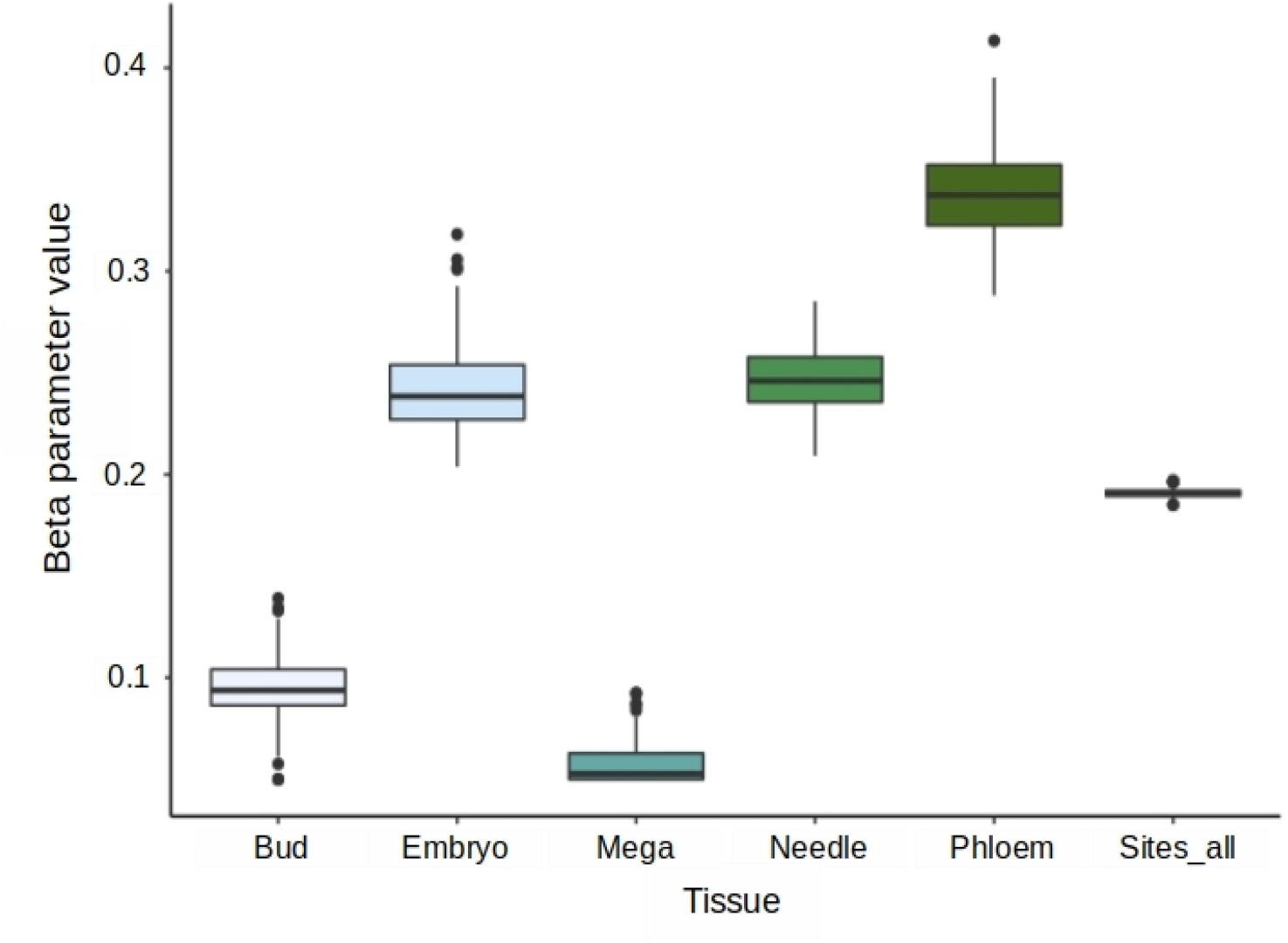
Distribution of the β parameter values obtained from 200 inferences of the DFE of 0-fold mutations for tissue-specific genes (tau score ≥ 0.8) showing the extreme distribution of the values observed in haploid megagametophyte genes.

Evolutionary expectations over gene expression breadth establish that narrowly expressed genes are under more relaxed selection. As haploid tissue-specific genes are narrowly expressed, this contrasting effect could conceal the patterns arising from haploid selection. Also, genes with higher levels of expression are under stronger selective constraints, which can confound the signal of purifying selection observed in megagametophyte tissue-specific genes. Hence, we looked at the extreme values of tau and the level of expression to see if our results were robust to these confounding factors. First, we restricted further the DFE estimations to genes with tau values above median. Our results show (Figure 4) that contrary to expectations over breadth of expression, highly tissue-specific genes in the megagametophyte show an even clearer signal of purifying selection compared to the other tissue-specific genes. Second, we evaluate if the strong signal of purifying selection we observed in megagametophyte tissue-specific genes was mainly driven by highly expressed genes or was independent of the level of expression. Hence, we subset the tissue-specific genes according to their level of expression, and we run a DFE inference on genes that range in expression from the lowest value up to the median value of expression. Our results show (Figure 5) that the signal of purifying selection in haploid-specific genes is not driven by megagametophyte specifically-expressed genes with higher levels of expression, but that also genes with low levels of expression show a strong signal of purifying selection.

**Figure 4.**
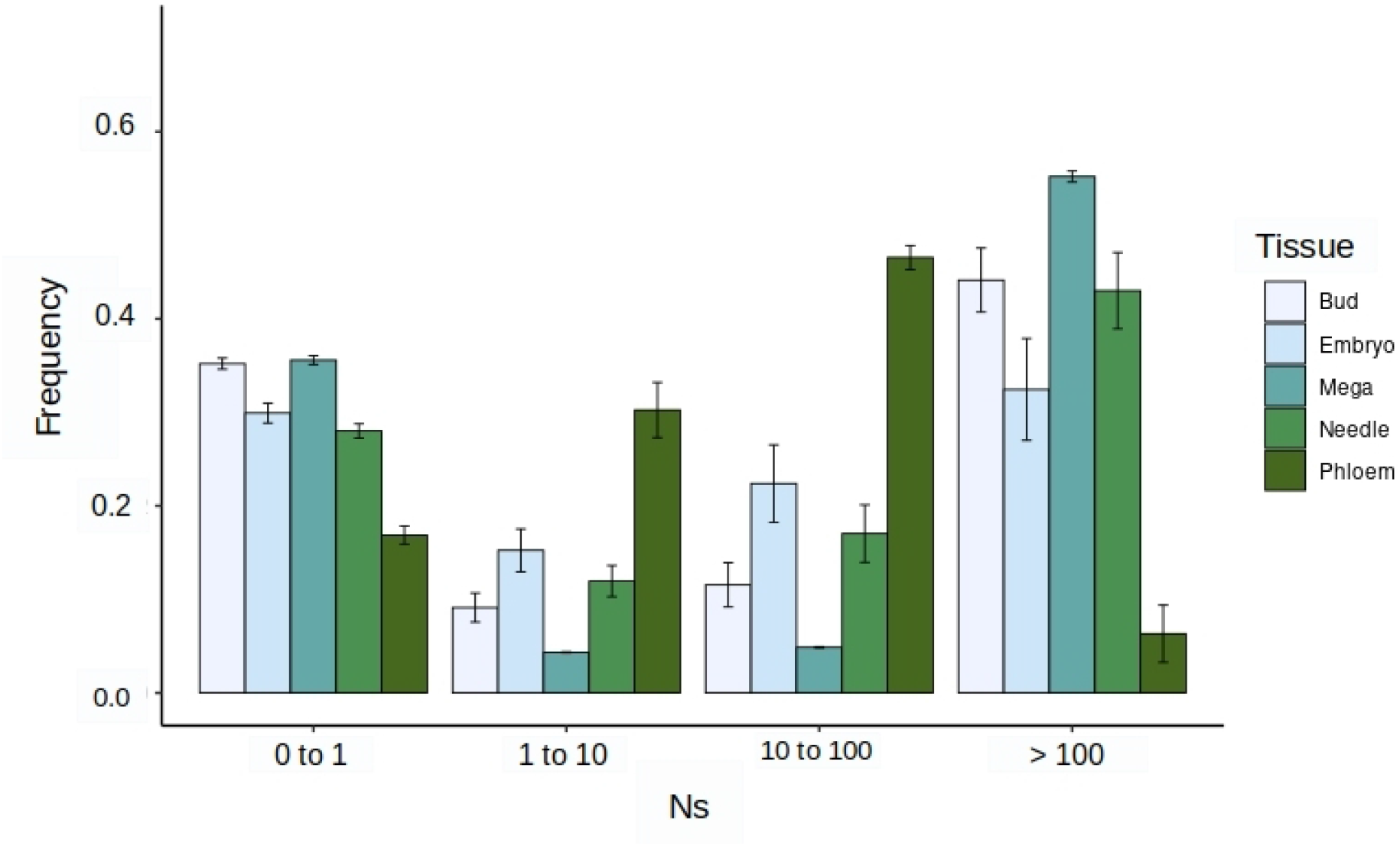
Distribution of fitness effect for highly tissue-specific genes (above median). Results show the DFE across 200 resampling of the fSFS.

**Figure 5.**
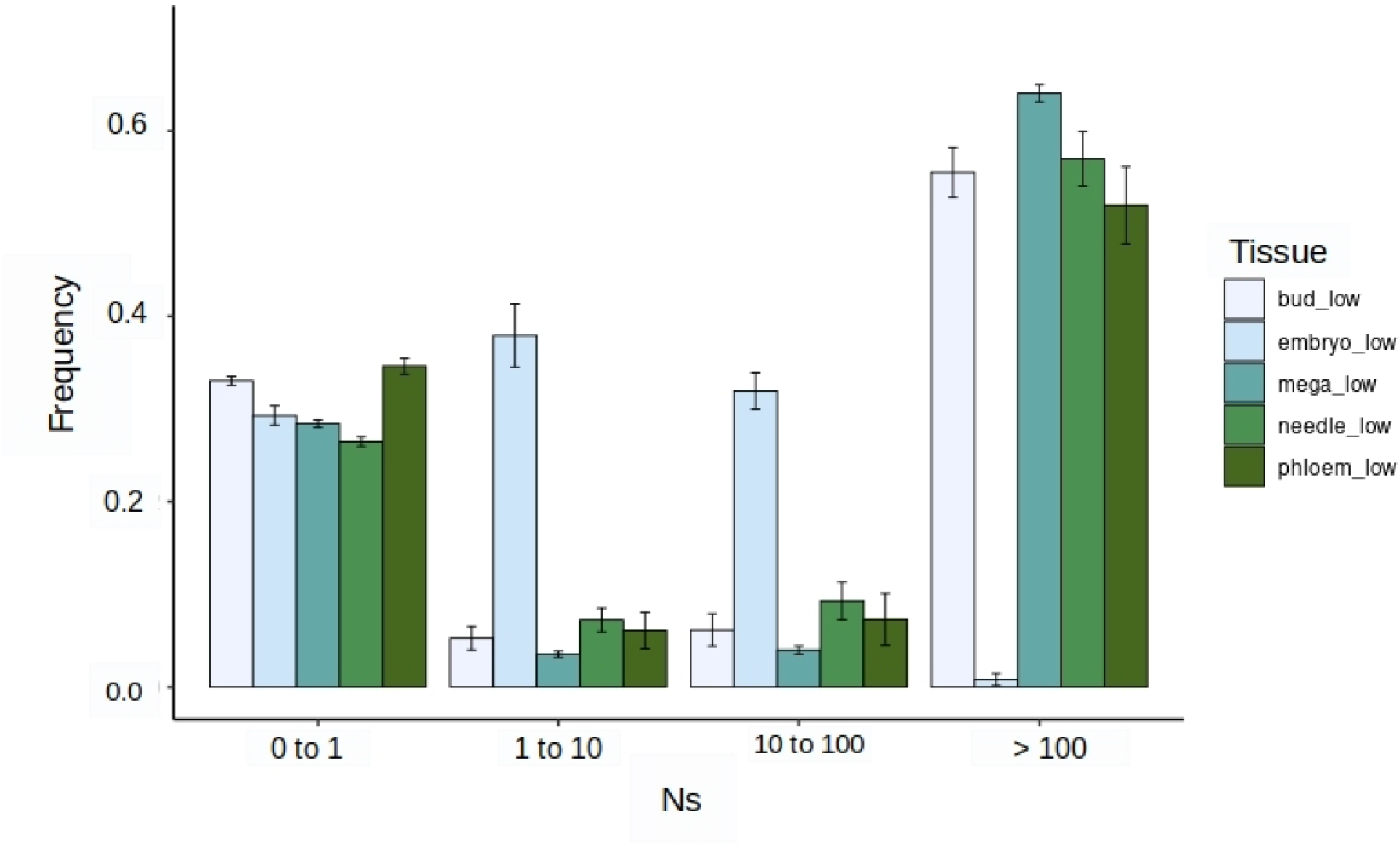
Distribution of fitness effect of tissue-specific genes with expression from the low to the median value of gene expression.

## Discussion

Here, for the first time, we present evidence of haploid selection on tissue-specific genes of a long-lived gymnosperm tree. Our findings demonstrate that purifying selection is stronger on genes with tissue-specific expression in haploid female-derived megagametophytic tissue compared to diploid tissue-specific genes. In comparison to other studies where the presence of haploid selection has been demonstrated (Chibalina and Filatov 2011; Arunkumar et al. 2013; Gutiérrez-Valencia et al. 2022) our results are free of the confounding effects of sexual competition. Szövényi et al. (2013) suggested that in *F. hygrometrica*, a non-angiosperm system also free of confounding effects of sexual competition, the low level and narrow breadth of expression of tissue-specific genes could explain the lack of significant differences in the amount of selective constraint between diploid and haploid tissue-specific expressed genes. Here, our results show that the strength of purifying selection varies across tissues with some diploid tissues, e.g. bud and needle, also having strong signals of selection. Evidence of selective constraint variation between genes expressed in different tissues has been reported earlier e.g., in a comparative study of 22 different tissue types (including reproductive and non-reproductive organs) of mouse (Kryuchkova-Mostacci and Robinson-Rechavi 2015). Similarly in plants, selective constraint differs between genes expressed in different female and male reproductive tissues, reflected as differences in the DFE (Gutiérrez-Valencia et al. 2022). However, we found that unlike diploid tissues, the signal of purifying selection of haploid tissue-specific genes in *P. sylvestris* was robust to further narrowing of the breadth of expression and to low level of expression - known determinants of evolutionary rate (Duret and Mouchiroud 2000). The signal of strong selection in megagametophyte is in line with both the theoretical expectation of strong selection at haploid stage (masking theory), and the known important function of megagametophyte as a nutrition-providing tissue for a germinating embryo (Vuosku et al. 2009).

The method used for the estimation of the DFE (Keightley and Eyre-Walker 2007) assumes additivity, where the effect on an allele does not depend on the other allele, although this assumption is often violated and ignored. For the purpose of comparative studies seeking qualitative differences in DFE among e.g., gene groups, the assumption of additivity is not a problem when there is no reason to believe that the distribution of dominance is different among the gene groups compared. In our study, dominance (*h*) has an essential role, as the masking is most efficient for new mutations with low dominance. For haploid-expressed genes, the assumption of additivity is insignificant, as all the alleles are exposed to selection independent of the other alleles and dominance level does not matter. However, for all diploid-expressed genes, dominance will have an effect on the DFE estimation. Higher dominance leads to more efficient purifying selection and vice versa. Thus, the differences we observe between haploid and diploid expressed genes DFE are not only reflecting their *Ns*, but also *Nhs* (Huber et al. 2018). Thus, some proportion of DFE differences are probably caused by masking of new mutations with *h* lower than 0.5, moving the *Ns* estimate closer to neutral/nearly neutral class (*Ns*<1) when expressed in the diploid stage.

Unlike the clear signal of purifying selection in haploid tissue-specific genes, we did not observe a striking difference in the levels of genetic diversity among tissue-specific genes expressed in the haploid and diploid stages. However, in effective outcrossers, such as *P. sylvestris*, with low levels of linkage disequilibrium the signal of background selection can be less evident due to the effects of recombination (Charlesworth et al. 1995). We neither observed a skew towards rare alleles in the fSFS of haploid tissue-specific genes, but instead observed excess of completely invariant sites, as would be expected under very strong selection. In diploid genomes recessive deleterious alleles can remain segregating in low frequency because they are rarely exposed to selection as homozygotes. This can be reflected in the SFS as a skewness toward rare alleles. However, in haploid-expressed genes even rare recessive alleles are exposed to selection, which may explain why polymorphisms of 0-fold sites are not enriched among rare allele classes in megagametophyte tissue-specific genes.

Haploid life-stages are common in nature, with green plants displaying great variation in the spatial and temporal extent of haploid stage, from dominantly haplontic liverworts to diplontic angiosperms (e.g., Mable and Otto 1998). Nevertheless, studies on the effects of selection tend to concentrate on the sporophyte-stage, especially in species with less conspicuous haploid-stage. In this study, we observed a strong signal of purifying selection in genes expressed in a reproductively important and relatively long-lived tissue: gymnosperm female gametophyte. Our results, as well as results in other systems such as Rumex (reference), show that genes expressed in haploid-stages can be exposed to effective selection. Selection over haploid stages and female tissues deserves more attention also because it can affect the evolutionary dynamics of the sporophyte-stage, depending on the amount of pleiotropy across sporophyte and gametophytes.

## Materials and methods

### Biological material and DNA extraction

We obtained seeds for nucleotide diversity-based analysis from 20 randomly-selected open-pollinated *P. sylvestris* trees from a single naturally-regenerated population at the Punkaharju Intensive Study Site Finland (https://www.evoltree.eu/resources/intensive-study-sites/sites/site/punkaharju) managed by Natural Resources Institute Finland (Luke). We induced germination by placing the seeds in Petri dishes with wet paper and incubating them overnight in dark at room temperature. We dissected the megagametophytes from the germinating seeds and extracted DNA with E.Z.N.A Plant DNA DS Kit (Omega BIO-TEK). We quantified the DNA concentration with NanoDrop ND-1000 (Thermo Fisher Scientific) and fragmented DNA by sonication with a Bioruptor UCD-200 (Diagenode) using two periods of 15 minutes and one period of 13 minutes, with periods consisting of cycles of 30 s on and 90 s off. We did a double-side size selection with AMPure XP beads (Beckman Coulter) targeting fragments between 300 to 350 bp. We then confirmed the fragment distribution on a 2100 Bioanalyzer (Agilent) using the Agilent DNA High Sensitivity chips.

### Library preparation and exome capture

We prepared DNA libraries using the Kapa HyperPrep (Roche) library kit according to manufacturer’s protocol and libraries were indexed with Kapa Single-Indexed Adapter kits A and B. Indexed libraries were enriched with 3 to 4 PCR cycles (depending on library concentration). We verified the quality and the length distribution of the libraries with 2100 Bioanalyzer and High Sensitivity chips (Agilent), and their concentration with Qubit HS dsDNA kit (Thermo Fisher). To obtain a reduced representation of the genome we used the PiSy UOULU exome capture design (Roche) described in Kesälahti R. unpublish data. We set two hybridization reactions each containing species-specific c0t-1 DNA, the exome capture baits, and 10 samples pooled equimolarly to a total amount of 1000 ng of DNA (Kesälahti R. unpublish data). Each hybridization reaction was incubated for 18 hours, followed by 14 PCR cycles for enrichment. The final pools were quantified using the Kapa Library Quant kit (Roche) according to manufacturer’s protocol in a LightCycler 480 (Roche). After quantification we pooled equimolarly the two hybridization reactions in a single sample. The sequencing was done in an Illumina NextSeq550 with 150 bp paired-end reads at the Biocenter Oulu Sequencing Centre.

### Mapping and variant call

Demultiplexed and adapter-removed reads of the 20 samples obtained from the sequencing facility were mapped to the *P. taeda* reference v2.01 (https://treegenesdb.org/FTP/Genomes/Pita/v2.01/) (Zimin et al.2017) using BWA (Li and Durbin 2009) with default parameters. SAM files were converted to BAM and sorted using Picard tools 2.21.4. We filled coordinates information in the bam files with samtools 1.9 (Li et al. 2009) fixmate, sorted the files by leftmost coordinates, and removed duplicates with samtools markdup. We added read groups with Picard tools, and indexed the final bam files with samtools index.

We did two joint variant calls with Freebayes v 1.3.1 (Garrison and Marth 2012). The first call (hereafter diploid call) was done with default parameters and a ploidy level of two. The objective of this variant call was to identify genomic regions with paralogy. As our samples originated from haploid tissue, we did not expect to observe any real heterozygosity. Hence, we used any observed heterozygosity as a proxy of different paralog copies mapping to the same genomic region (Tyrmi et al. 2020). We used BCFtools (Danecek et al. 2021) to identify SNPs positions with two or more heterozygous calls. We then identified the position surrounding the SNP in a 150 bp window with a custom R script (https://github.com/GenTree-h2020-eu/GenTree/blob/master/kastally/paralog_window_filtering/paralog_window_filtering.R). We consider all positions within the 150 bp windows as putative paralogous regions to be further removed.

For the second variant call (hereafter haploid call) we used Freebayes with the option ‘population model’ and a theta value of 0.005, ploidy level one, and report of invariant sites. We removed complex variants and sites with more than two alleles with BCFtools and indels with VCFtools (Danecek et al. 2011). We then removed the putative paralog regions identified in the diploid call with VCFtools. We filtered variant and invariant positions at genotype level for genotype quality (GQ) > 20, depth (DP) > 10 and maximum of 20% missing data with VCFtools. Then we used the vcffixup command from vcflib (Garrison et al. 2021) to update the allele number (AN) and allele count (AC) fields to reflect the final genotype counts on the VCFs.

### Identification of genes with tissue-specific expression

To identify genes with haploid and diploid tissue-specific expression and expression levels, we used data gene expression data of Cervantes et al. 2021 which had been mapped to *P. sylvestris* reference transcriptome (Ojeda et al. 2019, BioProject PRJNA531617). We linked the variant call data mapped to *P. taeda* to the gene expression data indicating the most likely homologous region between *P. sylvestris* transriptome and *P. taeda* reference (version 2.01) based on blast (Kesälahti R. unpublished data).

Next, we generated a bed file for all polymorphic and monomorphic positions in the vcf file with the function vcf2bed from BedOps v 2.4.38 (Neph et al. 2012). Then, we used BedTools (Quinlan and Hall 2010) intersect with the options -wa, -wb, and -loj, to link the positions in the vcf file to the information in Kesälahti R. unpublished data (supplementary data 1 and 2). We only retained positions that had a unique match in the *P. sylvestris* transcriptome.

For each gene, we obtained the tissue-specificity and gene expression level information across five tissues (megagametophyte, haploid, and four diploid tissues vegetative bud, embryo, needle, and phloem) from the TMM matrix reported in Cervantes et al. 2021. We considered genes with a tau index score of 0.8 and up as tissue-specific (Yanai et al. 2005).

### Structural annotation of variant and invariant positions

We used the NewAnnotateRef.py script (https://github.com/fabbyrob/science/blob/master/pileup_analyzers/NewAnnotateRef.py) and the *P. taeda* gtf file of v2.01 genome (https://treegenesdb.org/FTP/Genomes/Pita/v2.01/annotation/) to obtain the 0-fold and 4-fold positions of the *P. taeda* scaffolds where we had mapping information. After this, we used VCFtools with the –positions option and the obtain one VCF file with monomorphic and polymorphic positions at 0-fold and 4-fold sites (supplementary data 3 and 4).

### Genetic diversity estimates

We calculated π at 0-fold and 4-fold sites for tissue-specific genes (tau ≥ 0.8) with pixy (Korunes and Samuk 2021). We used the all-genes dataset as an input for intervals having a corresponding unique *P. sylvestris* transcript and --bypass_invariant_check as ‘no’ to include invariant sites. We kept only genes for which we have both 0-fold and 4-fold estimates available (supplementary data 5). Additionally, we retained only genes where the total number of sites (0-fold plus 4-fold) used for the estimates of diversity was 50 bp or larger. To obtain standard errors we used R to obtain a distribution of means by bootstraping 1000 times over the per gene π estimates (Pruim et al. 2017, Wickham et al. 2019, R Core Team 2020). Amount of genes used per tissue was 378 for bud, 284 for embryo, 332 for mega, 579 for needle, 387 for phloem, and 1960 for all-genes dataset.

### Site frequency spectrum and distribution of fitness effects (DFE)

We calculated the distribution of fitness effects for the five sets of tissue-specific genes and the all-sites dataset using dfe-alpha 2.16 (Keightley and Eyre-Walker 2007) using the fSFS for the 0-fold and 4-fold sites. To obtain the fSFSs we used the bait positions for each set of tissue-specific genes and generated two VCF files per each dataset, one for 0-fold and another for 4-fold. Since the amount of missing data varies across sites, we down-sampled without replacement 200 times the fSFS to five different sample sizes from 20% to 0% missing data (16, 17, 18, 19, and 20 alleles)

(https://github.com/H2020-GenTree/GenTree/tree/master/kastally/sfs_resampling). To account for positions monomorphic among *P. sylvestris* samples, but carrying a different allele than the *P. taeda* reference, we set all the AC (alternate count) position to zero for sites where AN (allele number) = AC. We then used the five downsampled fSFS as input for the estimation of the DFE with a 2-epoch model of inference. For the neutral (4-fold) sites we searched for the best N_2_ (population size after the change of Ne), t_2_ (duration of the epoch of population size change), and an initial t_2_ of 50 generations. For the selected sites (0-fold) mean effect of deleterious mutations (s) had an initial value of -0.1, and beta had an initial value of 0.5.

### Overlap of DFE inferences

To quantify the amount of overlap between the β parameter values of the bud tissue-specific genes and the megagametophyte tissue-specific genes, we generated a density distribution plot for the β parameter values obtain from the 200 inferences of the DFE (Supplementary Figure 1). Then, to determine the percentage of area overlapping we applied the procedure described at https://stackoverflow.com/questions/64882496/how-to-calculate-the-overlap-between-2-dataset-distribution. Colors for all figures included were obtain from R package Pacific North West Colors (Lawlor 2020).

## Acknowledgment

This work was supported by the Academy of Finland grants 287431, 293819 and 319313 to TP, Biocenter Oulu (SC), and Emil Aaltonen Foundation (RK). The authors acknowledge the CSC – IT Center for Science, Finland, for computational resources.

## Author Contributions

The study was designed by TP. All lab work was done by TK. Initial bioinformatic work was done by SC, RK and TM. Annotation and all analyses were done by SC. Writing of draft manuscript was done by SC. TP and HH contributed substantial comments to the manuscript and guidance of all analyses. All authors reviewed and commented on the manuscript.

## Data Availability

Raw read for the exome capture are deposit at NCBI BioProject PRJNA937910

## Supplementary material

All listed supplementary data are at Figshare project: https://figshare.com/s/494380953602ea5fe016

