## Supplementary Fig. 1 for "Strong purifying selection in haploid tissue-specific genes of Scots pine supports the masking theory"

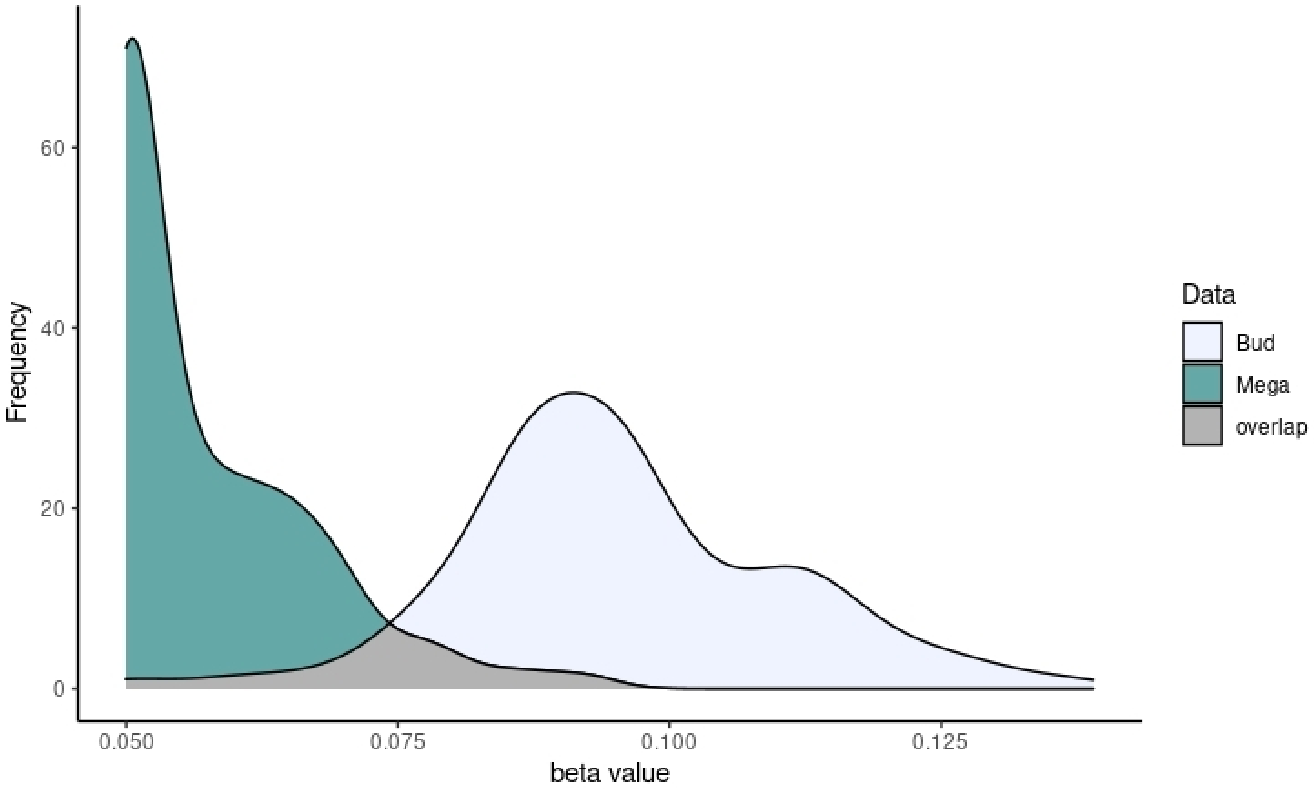


Supplementary figure 1. Density distribution of the β parameter values for the bud and megagametophyte tissue-specific gene sets and the all-sites showing the overlap area.
